# Natural variation in the *Arabidopsis AGO2* gene is associated with susceptibility to potato virus X

**DOI:** 10.1101/746628

**Authors:** Chantal Brosseau, Ayooluwa Adurogbangba, Charles Roussin-Léveillée, Zhenxing Zhao, Sébastien Biga, Peter Moffett

**Author notes:** **Corresponding author**: Peter Moffett, Département de Biologie, Université de Sherbrooke, 2500, Boulevard de l’Université, Sherbrooke, Québec, Canada, J1K 2R1.

## Abstract

RNA silencing functions as an anti-viral defence in plants through the action of DICER-like (DCL) and ARGONAUTE (AGO) proteins. However, there are few known examples of functional variation in RNA silencing components. The AGO2 protein is important for antiviral defense against multiple viruses and has been shown to be a major limiting factor to infection by potato virus X (PVX) of *Arabidopsis thaliana* but not *Nicotiana benthamiana*. We show that the AGO2 proteins from these two plants have differential activity against PVX, suggesting that variation in AGO2 is important in plant-virus interactions. Consistent with this, we find that the *Arabidopsis thaliana AGO2* gene shows a high incidence of polymorphisms between accessions, with evidence of selective pressure. AGO2 protein variants can be assigned to two groups, in near equal frequency, based on an amino acid change and small deletions in the protein N-terminus. Inoculation of a large number of *Arabidopsis* accessions shows strong correlation between these alleles and resistance or susceptibility to PVX. These observations were validated using genetic and transgenic complementation analysis, which showed that one type of AGO2 variant is specifically affected in its antiviral activity, without interfering with other AGO2-associated functions such as anti-bacterial resistance or DNA methylation. Our results demonstrate a novel type of genetically-encoded virus resistance and suggest that plant-virus interactions have influenced natural variation in RNA silencing components.

## Introduction

RNA silencing is a conserved gene regulatory mechanism that is also employ by plants to counteract virus infection^1^. Virus double-stranded RNA (dsRNAs), produced during the replication of RNA viruses, is recognized and cleaved into small-interfering RNAs (siRNAs) by dicer-like (DCL) proteins. These siRNAs are then incorporated into an RNA-silencing complex (RISC) which contains Argonaute (AGO) endoribonucelase proteins. These programmed complexes subsequently target any single-stranded (ssRNA) with sufficient complementarity for cleavage or translation inhibition^2^.

To counteract RNA silencing, plant viruses encode viral suppressors of RNA silencing (VSR), which have been shown to interfere with different steps of this mechanism^3^. The P25 protein, also known as TGB1, of potato virus X (PVX) is implicated in viral movement, in the formation of viral replication complexes (VRC), and functions as a VSR by inducing a destabilization of multiple AGO proteins^4–7^.

Among the ten AGO proteins encoded by *Arabidopsis thaliana*, AGO2 has been identified most frequently as having antiviral function and this, against multiple different viruses^1^. Indeed, AGO2 appears to be a major determinant of the inability of PVX to systemically infect *Arabidopsis* accession Col-0^8^ and an antiviral role for AGO2 is conserved in *Nicotiana benthamiana* ^9,10,11,12^. However, despite the presence of a functional AGO2, *N. benthamiana* is susceptible to many viruses, including PVX^13^. Although multiple studies have elucidated how AGO proteins carry out their basic biochemical functions^11,14^, less is known about why certain AGO proteins function more or less effectively against viruses. Likewise, natural variation in RNA silencing components, including AGO proteins, within and between species, has not been extensively studied.

Here, we show that AGO2 proteins from *A. thaliana* and *N. benthamiana* manifest differing activities against PVX, suggesting that inter-specific differences in AGO2 may contribute to differing outcomes of PVX infection in these species. Furthermore, we tested whether intra-specific differences in AGO2 might affect plant-virus interactions by taking advantage of the natural genetic variation of wild *Arabidopsis* accessions. We show the *AGO2* gene presents a high level of polymorphism and shows evidence of having been subject to selective pressure. Furthermore, unlike the commonly used *Arabidopsis* accession Col-0, 27 out of 63 accessions analysed were found to susceptible to PVX. Through genetic and transgenic analysis, we show that this susceptibility is determined by two polymorphisms found in the N-terminus of the AGO2 protein. Our results have uncovered a novel form of genetically-encoded virus resistance. Likewise, it suggests that natural variation in *AGO2* may be important for determining plant-virus interaction outcomes and that in turn these pressures may have shaped the RNA silencing machinery in ways similar to other defense mechanisms.

## Results

### AGO2 proteins from different genera display specific antiviral activity

To determine if differences in PVX susceptibility between *N. benthamiana* and *A. thaliana* might be determined in part by AGO2, we transiently expressed the two proteins with PVX-GFP. Consistent with our previous results^15^, transient expression of AtAGO2 in *N. benthamiana* resulted in a lower PVX-derived GFP accumulation (Fig. 1a, 1b). However, NbAGO2 had much less effect on PVX accumulation, as determined visually and by immunoblotting (Fig. 1a, 1b). Despite being expressed under the same strong promoter, the two AGO2 proteins did not accumulate at similar levels in this experiment (Fig. 1b, middle panel). Previous studies have shown that the Potexvirus VSR, P25, has the potential to compromise stability or accumulation of different AGO proteins^7,15,16^. To monitor whether the presence of P25 affects NbAGO2, we used a mutant version of PVX, PVX-GFPΔTGB, which lacks P25. In contrast to WT PVX, both AGO2 proteins significantly reduced the accumulation of virus-derived GFP from PVX-GFPΔTGB (Fig. 1c, 1d). We also noticed that AtAGO2 and NbAGO2 accumulated at similar levels when co-expressed with PVXΔTGB (Fig. 1d), suggesting that P25 may affect the two different AGO2 proteins differently. Consistent with this, expression of AGO2 proteins with P25 alone reduced NbAGO2 accumulation, but not AtAGO2 (Fig. 1e). Taken together, these results suggest that P25 VSR activity is at least partially responsible for the differential efficiency in targeting PVX observed between these two AGO2 proteins.

**Figure 1.**
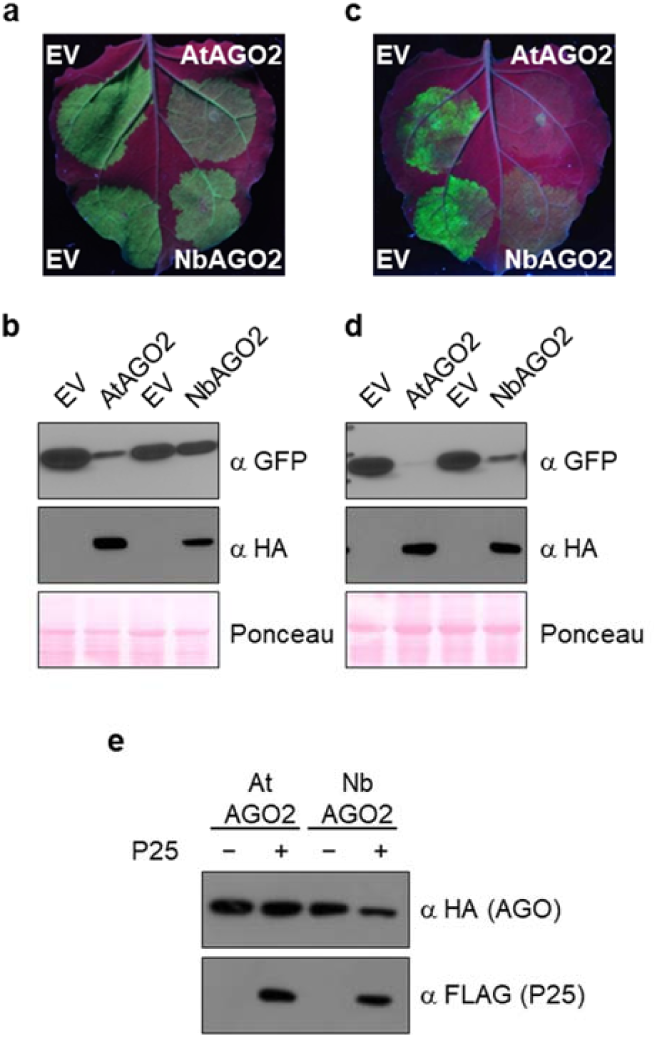
AtAGO2, but not NbAGO2, shows anti-viral activity against PVX. **a** and **c**, *N. benthamiana* leaves were agroinfiltrated with PVX-GFP WT **a** or ΔTGB **c** along with 35S:HA-AtAGO2, 35S:HA-NbAGO2 or empty vector (EV). Leaves were photographed under UV illumination at 4 days post infiltration (dpi). **b** and **d**, Total protein extracts were prepared from *N. benthamiana* leaves agroinfiltrated as in **a** and **c** at 4 dpi and subjected to SDS-PAGE, followed by anti-GFP immunoblotting (top panel). HA-tagged AGO proteins were immunoprecipitated from the same extracts and subjected to anti-HA immunoblotting (middle panel). Ponceau staining (bottom panel) of the same extracts is shown to demonstrate equal loading. **e**, HA-tagged AGO proteins were co-expressed by agroinfiltration in *N. benthamiana* leaves with either FLAG-tagged P25 or with empty vector (EV). Total proteins were extracted and subjected to anti-FLAG immunoblotting (bottom panel). HA-tagged AGO proteins were immunoprecipitated and subjected to anti-HA immunoblotting (top panel).

### The *Arabidopsis AGO2* gene displays a high degree of polymorphism

The NbAGO2 and AtAGO2 proteins are only 50% identical^17^, which may explain their being differentially affected by PVX P25, but makes it difficult to identify specific residues important for anti-PVX activity. To evaluate whether the *AGO2* gene shows differences within a species, we analyzed the coding sequences of *AGO1* (At1g48410) and *AGO2* (At1g31280) from 80 *Arabidopsis* accessions, representing eight Eurasian geographic regions^18^, as obtained from the 1001 genomes project (The 1001 Genomes Consortium, 2016) and by resequencing the *AGO2* gene from some accessions (Supplementary Table 1). Upon analysis, we observed a high level of single-nucleotide polymorphisms (SNP) throughout the *AGO2* coding sequence (Fig. 2a and Supplementary Table 1). Synonymous SNPs are six times more frequent in the *AGO2* coding region compared to *AGO1*, while non-synonymous SNPs are more than fifty times more frequent (Supplementary Table 1 and 2; Fig. 2a), suggesting that *AGO2* has been subjected to selection pressure.

**Figure 2.**
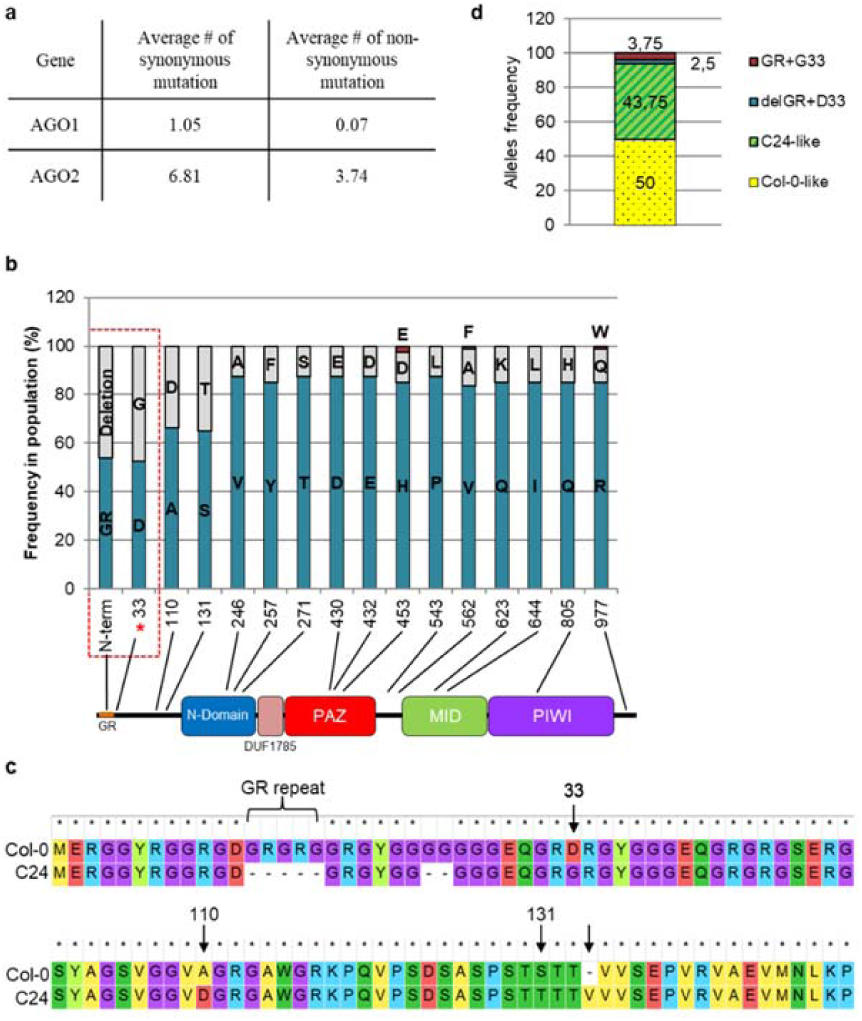
Residue 33 of *Arabidopsis AGO2* has undergone positive selection in natural populations. **a**, Variability observed in *AGO1* and *AGO2* sequences in 80 Eurasian *Arabidopsis* accessions in comparison to the Col-0 accession. **b**, Chart showing AGO2 residues having a high frequency of non-synonymous polymorphisms in the 80 accessions. A red dotted rectangle identifies polymorphisms frequently co-occurring in AGO2. An asterisk indicates an amino acid variant found to be subjected to positive selection pressure supported by fixed-effects likelihood (FEL)^57^ with a *p* value < 0.1 using DataMonkey.org website. Different domains of AGO2 are schematically represented by rectangles (not to scale) under the chart. Blue: N-terminal, red: PAZ, green: MID and purple: PIWI. **c**, Alignment of the N-terminal region of AGO2 protein of Col-0 and C24 AGO2 alleles. **d**, Graph representing allele frequencies in 80 different Eurasian accessions based on polymorphisms found in GR motif and residue 33 of AGO2 N-terminal region.

Selective pressures on AGO2 sequences were evaluated by analysing the ratio of nonsynonymous (*dN*) to synonymous substitution rates per site (d*S*). Site-by-site analysis indicated that residue 33 (Col-0 allele) showed the strongest signal of having undergone positive selection pressure (Fig. 2b). In the 80 accessions analyzed, only two different amino acids are found at position 33, namely an aspartic acid or a glycine (Supplementary Table 1, Supplementary Fig. 1a). The Col-0 accession encodes an aspartic acid at position 33 whereas many other accessions, including the commonly used C24 accession, encodes a glycine at the equivalent position (Fig. 2c). We also observed that multiple *AGO2* alleles possessed short deletions (compared to Col-0) resulting in deletions of 2 to 13 amino acids, in the region of the protein N-terminal to residue 33, which encodes a number of GR repeats (Fig. 2b), as is seen in the C24 accession (Fig. 2c, Supplementary Fig. 2). Interestingly, these deletions are almost always correlated with the presence of a glycine at residue 33 (Fig. 2b, Supplementary Fig. 1a). Given this strong correlation, we refer to *AGO2* alleles encoding 33D as Col-0-like and those encoding 33G plus a GR deletion as C24-like, although C24 *AGO2* possesses 2 additional SNPs, resulting in amino acid changes A110D and T131S, as well as an insertion of a valine at residue 134.

Although multiple *AGO2* alleles showed additional indels and SNPs, none of these individual differences were present at high prevalence. In the set of 80 Eurasian *Arabidopsis* accessions investigated, Col-0-like and C24-like alleles are present at a frequency of 50% and 43.75%, respectively (Fig. 2d) and both alleles are found in all eight sub-populations (Supplementary Fig. 1a). Likewise, a phylogenetic analysis of *AGO2* sequences failed to show geographic clustering, although such analysis was largely inconclusive, due to low bootstrap values (Supplementary Fig. 3). Only three variants with a full GR motif plus G33 and only two variants with a GR deletion plus D33 were identified, which we refer to collectively as rare *AGO2* alleles (Fig. 2d and Supplementary Fig. 1a). Interestingly, the presence of an aspartic acid at the equivalent of residue 33 of AGO2 appears to be the exception in the *Brassicaceae* family, with other species investigated encoding a glycine at the equivalent residue (Supplementary Fig. 1b).

### C24-like *AGO2* alleles are strongly associated with systemic infection by PVX in *Arabidopsis*

To test whether sequence variation observed in AGO2 might influence antiviral activity, we inoculated 63 accessions from the eight different populations with PVX. Susceptibility or resistance was scored based on the detection of PVX CP in systemic tissues (i.e. non-inoculated upper leaves) by immune blotting. We observed that, unlike Col-0, multiple accessions were susceptible to PVX and that resistance or susceptibility did not correlate with geographic origin (Fig. 3a; Supplementary Fig. 2 and 3; Supplementary Table 1). However, we found that 27 out of 30 tested accessions possessing a Col-0-like *AGO2* were resistant to PVX (Fig. 3; Supplementary Table 1). Conversely, 24 out of 31 tested accessions having either a C24-like or a rare *AGO2* allele were susceptible to PVX (Fig. 3; Supplementary Table 1). The *JAX1* gene has been shown previously to confer broad-spectrum resistance to potexviruses and its presence varies between *Arabidopsis* accessions^19^. We thus determined the *JAX1* status (+/-) in all accessions tested (Fig. 3; Supplementary Table 1). After doing so, we found that all PVX-resistant accessions with a C24-like *AGO2* allele, except Copac-1, also possessed a functional *JAX1* gene (Fig. 3). Together, these results indicate that variability in *AGO2* sequence is strongly associated with susceptibility to PVX in *Arabidopsis*.

**Figure 3.**
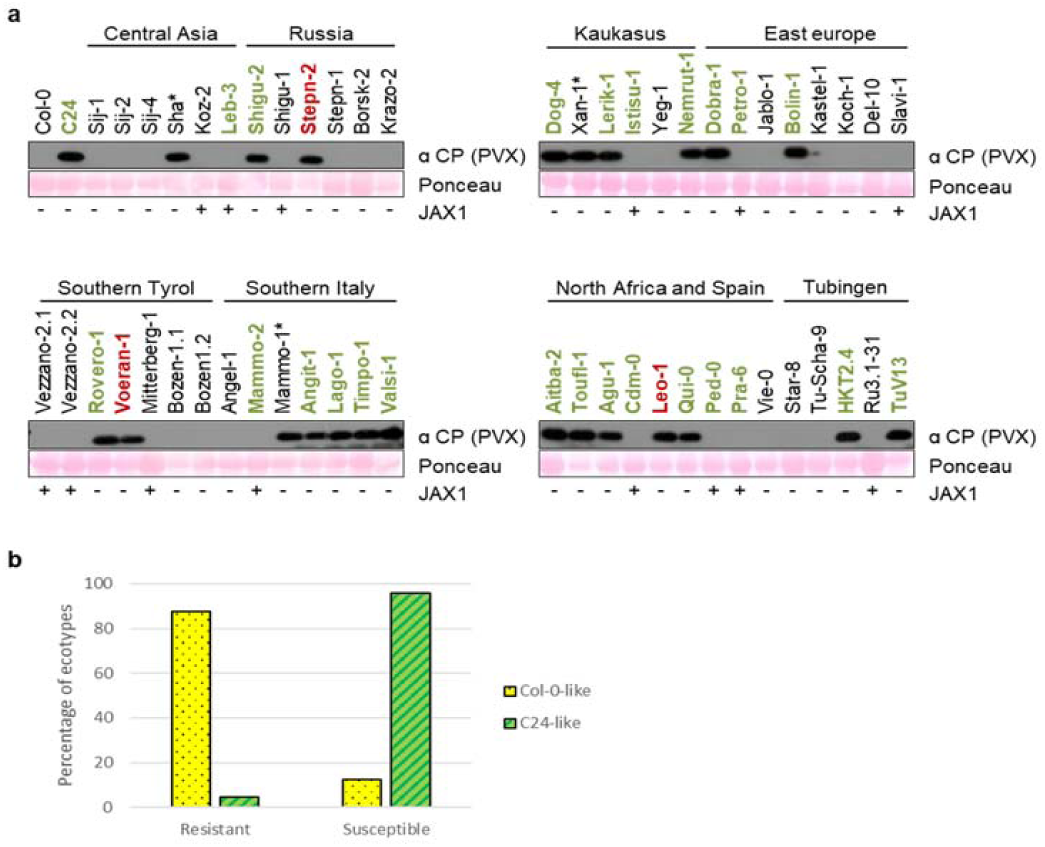
Natural variation in the N-terminus of AGO2 correlates with susceptibility to PVX. **a**, Different *Arabidopsis* accessions were inoculated with PVX, as indicated, categorized by geographic region of origin. At 21 dpi, total protein extracts were prepared from systemic leaves and subjected to SDS-PAGE followed by anti-PVX CP immune blotting. Ponceau staining (bottom panel) of the same extracts is shown to demonstrate equal loading. Accession names are colored according to their AGO2 allele: Col-0-like are black, C24-like are bold green and rare alleles are bold red. Asterisks indicate accessions that do not fit the expected correlation. Accessions were also genotyped *in silico* for the presence or absence (+/-) of a functional *JAX1* allele. Note that less protein sample was loaded into wells for C24, Stepn-2 and Bolin-1 due to the strong accumulation of PVX in these accessions. **b**, Compilation of results obtained for all JAX (-) accessions tested as in **a**, wherein the percentage of ecotypes are grouped by resistance or susceptibility to PVX and by the presence of either a Col-0-like or C24-like allele. The Pearson’s r-coefficient is 0.9958207 and falls within a 95% confidence interval.

### Validation of the effect of different AGO2 alleles and susceptibility to PVX in reciprocal inbred lines

Although we observed a strong association between *AGO2* alleles and PVX susceptibility, we cannot rule out that polymorphisms in other genes might contribute to the observed phenotypes due to the high level of genetic diversity between the accessions tested^18^. We thus took advantage of previously described reciprocal introgression lines (RILs). These lines were derived by crossing Col-0 and C24, followed by iterative backcrossing to the parental genotypes and selfing, resulting in lines with a majority of one parental genotype, with small genomic regions derived from the other^20^. From this collection, we selected RILs wherein the *AGO2* alleles were exchanged between accessions, as well as control lines wherein genomic regions adjacent to, but not including *AGO2*, were exchanged (Fig. 4a and Supplementary Table 3). RILs used in this study are depicted in Figure 4a and the origin of the *AGO2* allele was verified for each line by PCR (Supplementary Fig. 4a). We then assessed these lines for susceptibility to PVX. In agreement with our previous report^8^, no PVX was detected in the systemic tissues of Col-0, but was detected in more than half of the *ago2-1* mutant plants tested (Fig. 4b and Supplementary Fig. 4b). Similarly, PVX was not detected in systemic tissues of control RILs N35, N62 and N66, which have both Col-0 background and the Col-0 *AGO2* allele. However, PVX was detected in systemic leaves of more than half of infected plants from N37 and N55 lines wherein the C24 *AGO2* allele has been introgressed into a Col-0 background (Fig. 4b). Conversely, introgression of the Col-0-like *AGO2* allele into the C24 background (RIL M39) resulted in resistance to PVX similar to Col-0 plants (Fig. 4b and Supplementary Fig. 4b). A Pearson’s r-coefficient test showed a statistically significant positive correlation between the *AGO2* allele of the plant and its accumulation of PVX in systemic leaves (Fig. 4c).

**Figure 4.**
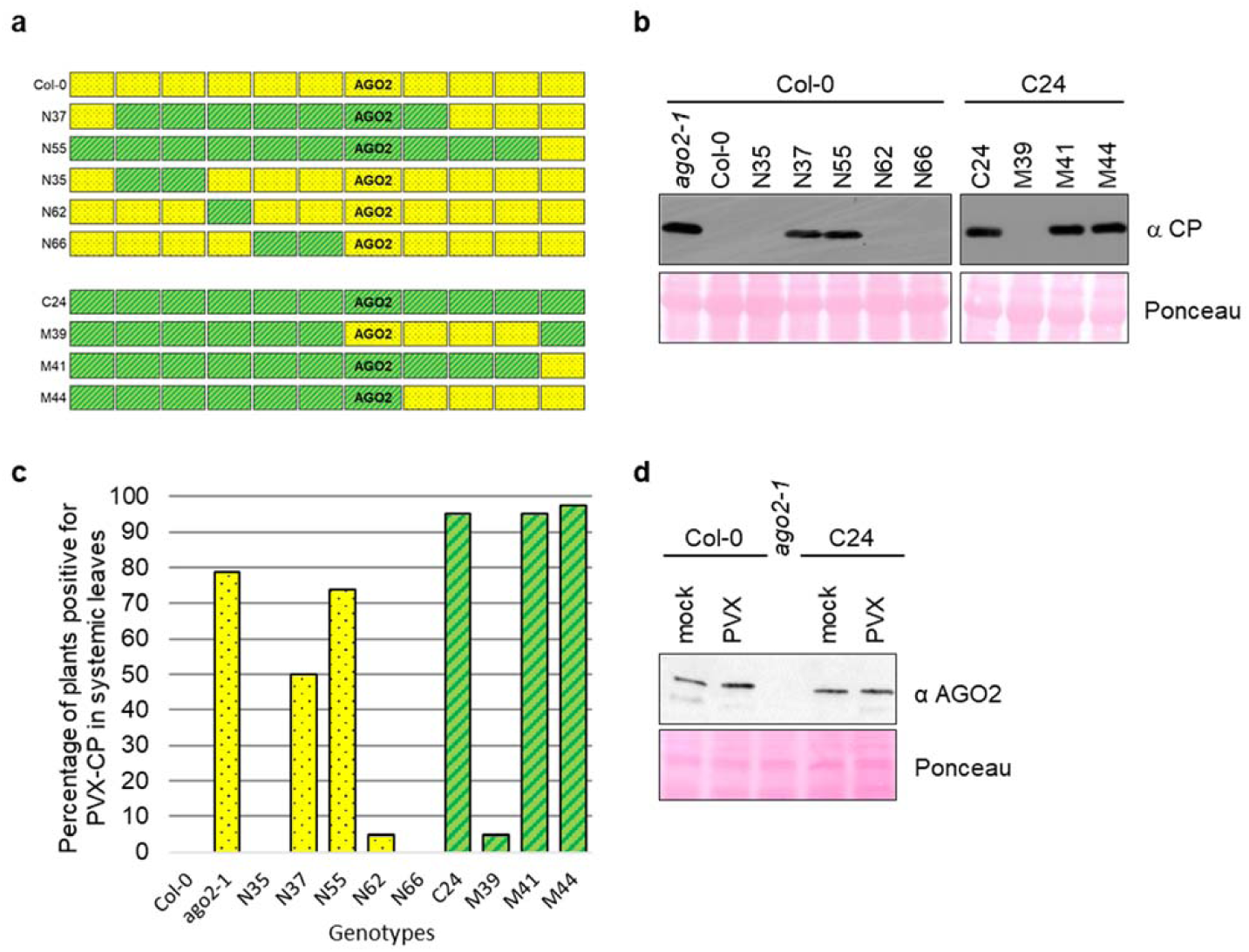
Exchange of *AGO2* alleles between Col-0 and C24 changes susceptibility to PVX. **a**, Schematic representation of the region of chromosome I between nt 174607 and nt 22286233 in recombinant intogression lines (RILs) from a cross between Col-0 and C24 (Törjék et *al*., 2008). N lines have a Col-0 background (yellow) throughout the genome except for small regions of chromosome I substituted with the corresponding region from C24 (green). M lines have a C24 background (green) with Col-0 substitutions (yellow). **b**, RILs, as depicted in **a**, were inoculated with PVX. At 21 dpi, total protein extracts from systemic leaves were prepared and subjected to SDS-PAGE followed by anti-PVX CP immune blotting (top panel). Ponceau staining (bottom panel) of the same extracts is shown to demonstrate equal loading. Representative results from 42 replicates are shown. **c**, Compilation of results obtained for all replicates tested as in **b** (related to Supplementary Fig. 2b). The Pearson’s rcoefficient is 0.6140612 and falls within a 95% confidence interval. **d**, Col-0, WT or *ago2-1*, and C24 plants were inoculated with PVX. At 21 dpi, total protein extracts from systemic leaves were prepared and subjected to SDS-PAGE followed by anti-AGO2 (Agrisera antibody) immune blotting (top panel). Ponceau staining (bottom panel) of the same extracts is shown to demonstrate equal loading.

The use of RILs between Col-0 and C24 accessions allows us to test the involvement of AGO2 alleles in virus resistance in the absence of potentially confounding effects of transgenes. At the same time however, several genes in this region of chromosome I display polymorphisms including *AGO3*. Phylogenetically, *AGO3* is the closest homologue to *AGO2* however, except against BaMV, *AGO3* has not been found to be involved in virus resistance^8,15,16,21,22^, but rather appears to play a role in DNA methylation^23^. Moreover, although C24 *AGO3* possesses some polymorphisms, *in silico* analysis revealed that these polymorphisms are also found in resistant accessions (Supplementary Table 3) precluding its implication in PVX susceptibility phenotype.

Exogenous application of the phytohormone salicylic acid has been shown to compromise PVX accumulation in *N. benthamiana*^24^ and *ICS1*, a key salicylic acid (SA) biosynthesis gene, is significantly up-regulated in C24 compared to Col-0^25^. However, quantification of SA in different RILs showed that introgression of the Col-0 genomic region containing the *AGO2* gene into the C24 background, or the opposite exchange, does not significantly change SA accumulation in these lines, relative to the parental genotypes (Supplementary Fig. 4c) thus precluding a role for SA in the observed phenotypes. Moreover, the differential susceptibility cannot be attributed to differential AGO2 expression as both accessions display similar AGO2 protein accumulation in systemic leaves upon inoculation with PVX (Fig. 4d). These results further validate the involvement of AGO2 polymorphism as a major determinant of resistance of to PVX in *Arabidopsis*.

### C24 *AGO2* is not a null allele

In *Arabidopsis*, AGO2 has also been implicated in antibacterial defense responses, presumably through its binding to endogenous miRNAs, as well as in the methylation of some DNA loci^26,27^. Aside from the differences found in the N-terminus outlined above, the C24 AGO2 protein does not differ from Col-0 in any of the well-characterized functional AGO domains. To determine whether C24 AGO2 is still efficient in non virus-related functions, we inoculated RILs with virulent *Pseudomonas syringae* pv. tomato (Pto). In Col-0 background lines in which C24 *AGO2* has been introgressed, namely N37 and N55, Pto grew at titres similar to WT and at significantly lower titers compared to *ago2-1* (Col-0 background) mutant plants (Fig. 5a). Likewise, introgression of Col-0 *AGO2* into C24 had no significant effect on bacterial growth compared to WT C24 (Fig. 5a). This suggests that C24 *AGO2* polymorphisms do not compromise the function of this protein in response to bacterial infection. Furthermore, we observed that Pst infection induces *AGO2* expression at similar level in both accessions (Fig. 5b), consistent with previous reports^26^.

**Figure 5.**
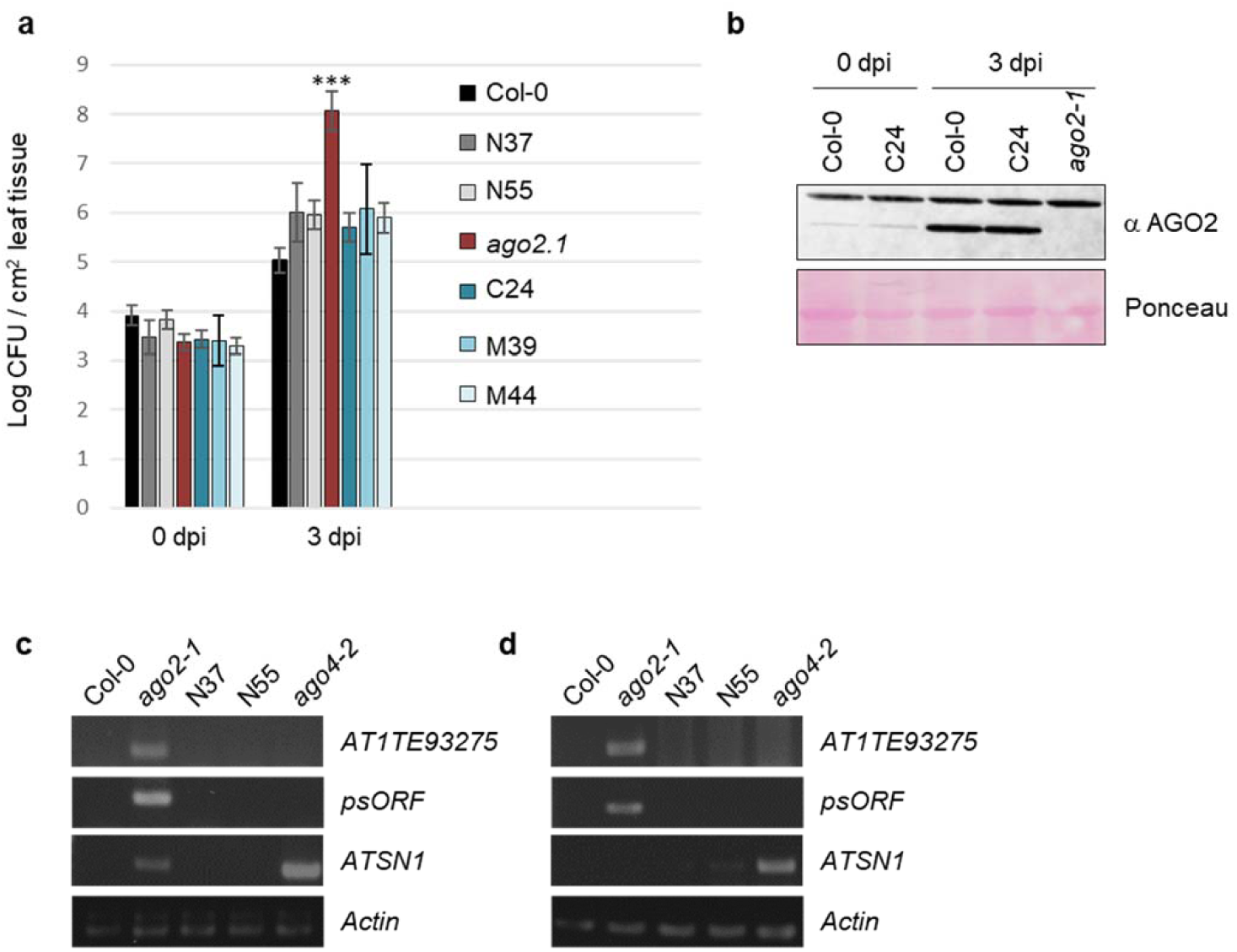
C24 AGO2 retains anti-bacterial and methylation-related functions. **a**, The indicated RILs with Col-0 and C24 backgrounds were infected with virulent Pst. Bacteria were counted at 0 and 3 dpi. Error bars indicate SEM from three biological replicates. Asterisks indicates statistically significant differences (student t test) at a P-value < 0.005. **b**, Total protein extracts were prepared from *Arabidopsis* leaves inoculated as in (A) at 0 and 3 dpi and subjected to SDS-PAGE, followed by anti-AGO2 (Abiocode) immunoblotting (top panel). Ponceau staining (bottom panel) of the same extracts is shown to demonstrate equal loading. **c**, Genomic DNA from the indicated genotypes was isolated and subjected to digestion with McrBC and subjected to PCR with primers to the indicated loci. **d**, RNA from the indicated genotypes was isolated and subjected to RT-PCR using primers to the indicated loci.

A previous study has reported that DNA methylation and gene silencing of the psORF and AT1TE93275 transposable elements are compromised in *ago2* mutant plants^27^. To examine whether C24 AGO2 is still functional in this respect, DNA methylation status and expression of AT1TE93275 and psORF were verified in two Col-0 lines containing the C24 *AGO2* allele. Genomic DNA from Col-0, *ago2-1*, N37, N55 and *ago4-2* plants was isolated and digested with the methylation dependent restriction enzyme McrBC. As shown in Figure 5c, the *ago2-1* mutant shows amplification of psORF and AT1TE93275 after McrBC digestion, whereas the N37 and N55 lines do not, while all lines show similar amplification of the actin gene (Fig. 5c). As expected, a similar test showed greater amplification of ATSN1, a signature locus for AGO4-dependent methylation, in the *ago4* mutant (Fig. 5c). At the same time, RT-PCR analysis showed that psORF and AT1TE93275 could be amplified only from RNA extracted from the *ago2-1* mutant and ATSN1 showed significant amplification only in the *ago4-2* mutant (Fig. 5d). These results are highly consistent with previous reports^27^ and indicate that both the Col-0 and the C24 AGO2 proteins are functional for AGO2-dependent methylation and its associated repression of a transposable element. Taken together, these results suggest that a complete GR motif and D33 of the Col-0 AGO2 protein are required for optimal antiviral defense but appear to be dispensable for regulating endogenous transcripts and methylation-related functions.

### C24 AGO2 shows decreased antiviral activity against PVX compared to Col-0 AGO2

Manual alignment of NbAGO2 and AtAGO2 proteins shows that NbAGO2 encodes a glycine at the position equivalent to G33 of C24-AGO2 (Supplementary Fig. 5). Because NbAGO2 was found to be efficiently antiviral only against a VSR-deficient version of PVX (Fig. 1), we verified whether it behaved similarly to C24 AGO2 in a transient overexpression assay. Indeed, in transient assays, C24 AGO2 is as efficient as Col-0 AGO2 at restricting PVX-GFPΔTGB but is somewhat less efficient against WT PVX-GFP (Supplementary Fig. 6a, 6b, 6c, 6d). This effect was also seen with other Col-0-like and C24-like AGO2 alleles, Yeg-1 and Bolin-1, respectively (Supplementary Fig. 6e, 6f, 6g, 6h). The difference between 35S-driven transient expression C24 vs. Col-0 AGO2 is not as dramatic as the difference seen with NbAGO2. However, expressing the different AGO2 variants under the native NbAGO2 promoter enhance the differential activity of these proteins against WT PVX (Fig. 6a, 6b) as well as against PlAMV-GFP, another Potexvirus (Supplementary Fig. 7a, 7b).

**Figure 6.**
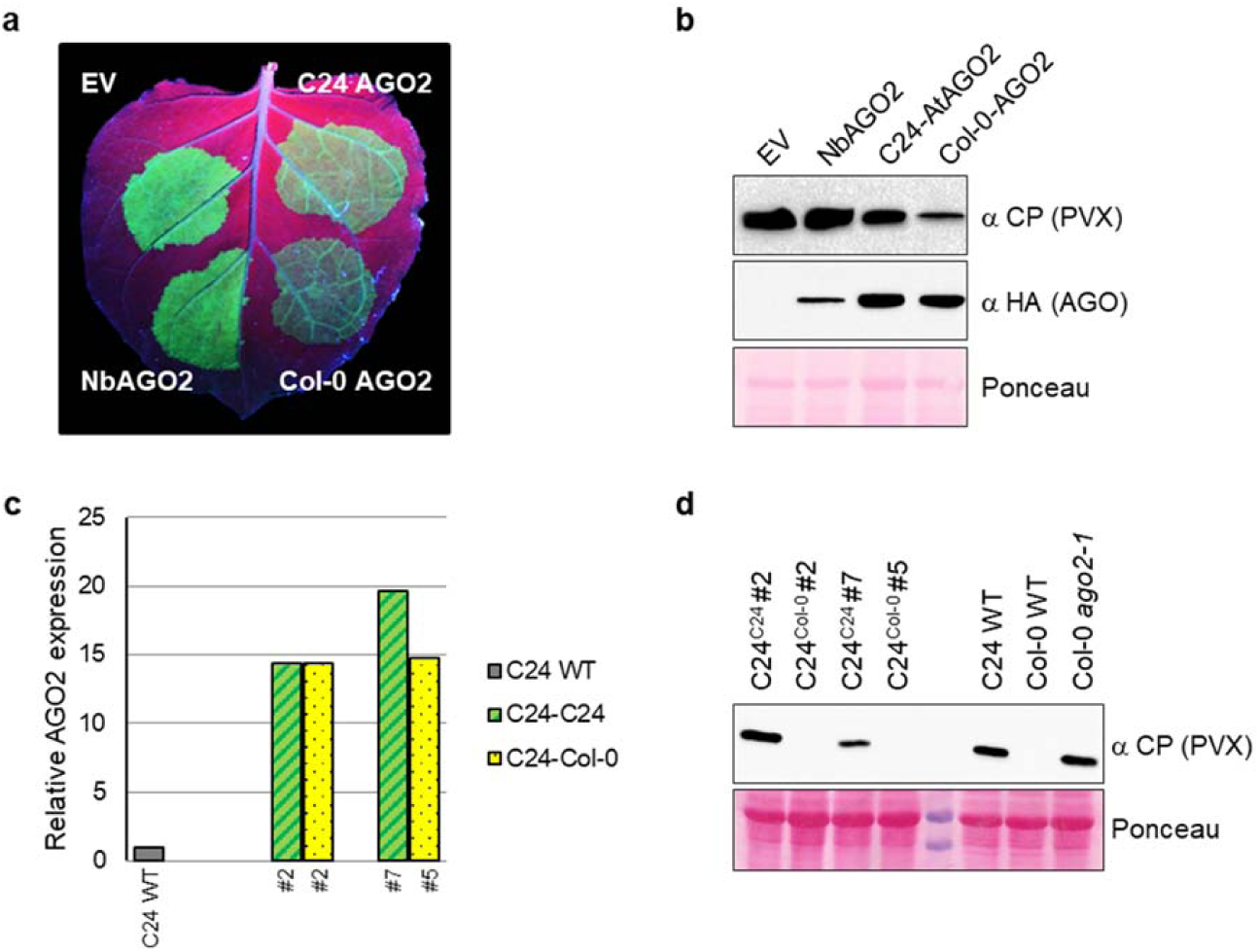
Polymorphisms found in C24 *AGO2* affect its antiviral activity in *Arabidopsis*. **a**, *N. benthamiana* leaves were agroinfiltrated with PVX-GFP, along with pNbAGO2:HA-NbAGO2, pNbAGO2:HA-AtCol-0-AGO2, pNbAGO2:HA-AtC24-AGO2 or empty vector (EV). Leaves were photographed under UV illumination at 4 dpi. **b**, Total protein extracts were prepared from *N. benthamiana* leaves agroinfiltrated as in **a** at 4 dpi and subjected to SDS-PAGE, followed by anti-PVX CP (top panel) or anti-HA (middle panel) immune blotting. Ponceau staining (bottom panel) of the same extracts is shown to demonstrate equal loading. **c** and **d**, WT C24, as well as different lines (#2, 5, 2 and 7) of transgenic plants expressing the indicated transgene, were inoculated with PVX. At 21 dpi, systemic leaves were harvested for qRT-PCR analysis **c** and immune blotting **d**. **c**, Total RNA was extracted from systemic leaves and subjected to qRT-PCR analysis to determine relative expression levels of total *AGO2*. Relative expression of *AGO2* in transgenic lines, C24^C24^ and C24^Col-0^, is normalized to the relative expression of *AGO2* in WT C24. **d**, Total protein extracts from systemic leaves were prepared and subjected to SDS-PAGE followed by anti-PVX CP immune blotting (top panel). Ponceau staining (bottom panel) of the same extracts is shown to demonstrate equal loading.

To verify whether this differential antiviral activity between Col-0-AGO2 and C24-AGO2 is biologically relevant in whole plants, we created C24 transgenic lines expressing Col-0-AGO2 or C24-AGO2 under the control of the 35S promoter. *AGO2* transgene expression was then monitored by qRT-PCR as, despite being under the control of a strong promoter, both proteins were undetectable by immune blotting analysis (data not shown). Lines with similar *AGO2* expression levels (C24^C24^#2 and C24^Col-0^#2) were analyzed by immune blotting to verify their susceptibility to PVX (Fig. 6c, 6d). At 21 dpi, PVX was not detected in systemic leaves of C24^Col-0^ lines, whereas C24^C24^#2 showed similar accumulation of PVX compared to that of C24 WT. In addition, the C24^C24^#7 line, expressing almost 20X more AGO2 than in C24 WT, remains susceptible to PVX as determined by immunoblotting (Fig. 6c, 6d). These results suggest that polymorphisms found in C24-AGO2 compromised its antiviral activity, and that AGO2 expression levels are not responsible for these differences. Combined with the RILs analysis, these results validate our conclusion that differences in AGO2 proteins are a major genetic determinant in resistance/susceptibility phenotypes observed in different *Arabidopsis* accessions.

## Discussion

### Susceptibility to PVX is common in *Arabidopsis thaliana*

Although the widely used *Arabidopsis* Col-0 accession is resistant to systemic PVX infection, we find that a large number of natural accessions are susceptible (Fig. 3). There are many potential barriers to compatible interactions between plants and viruses. However, our findings are consistent with studies showing that *Arabidopsis* encodes the necessary components to support Potexvirus replication^8,28,29^ and that abrogation of the RNA silencing machinery is sufficient to render Col-0 an effective host for PVX^8,15,19,30^. Combined with our findings that differential effectiveness of versions of AGO2 from different species and accessions, this suggests that natural variation in RNA silencing components may play important roles in host range determination and in ecological dynamics of plant-virus interactions.

### High prevalence of polymorphisms in the *AGO2* coding sequence

Surveys of *Arabidopsis* accessions have revealed extensive natural allelic variation, comprising SNPs as well as indels^31–33^. These naturally occurring variations are often associated with resistance to various biotic factors, particularly at disease resistant (R) gene loci encoding NLR proteins^34–36^. The *AGO2* coding region has 50 times more non-synonymous SNPs than the *AGO1* coding region (Fig. 2a; Supplementary Table 1 and 2). This suggests that AGO2 has been subjected to strong selective pressure for diversification. Such selection would be consistent with a study in *Drosophila*, showing that genes related to antiviral RNAi evolve at a significantly faster rate than paralogous genes implicated in housekeeping RNAi functions (miRNA pathway)^37^. Likewise, it has been proposed that viruses have driven close to 30% of all adaptive amino acid changes in mammals, making them dominant drivers of protein adaptation^38^. Importantly, in our analysis, we found no polymorphisms in residues predicted to be important for sRNA loading and maturation, RNA binding, AGO hook or catalytic functions^11^. Although variation has been observed throughout the coding sequence of *AGO2*, the only residue found to be under positive selection was in the extreme N-terminus, outside of conserved catalytic domains (Fig. 2), similar to what was observed in mammal RNAi-related proteins^37^. Consistent with the retention of core AGO function (Fig. 5) of C24 AGO2, this suggests that polymorphisms may have been selected for due to interactions with viruses. Indeed, all plant viruses are thought to encode at least one VSR, some of which interact directly with RNAi proteins, including AGOs, and thus host antiviral RNAi components must rapidly adapt to win the molecular arms race against viruses^37,39^. At the same time, VSRs must evolve to overcome the RNA silencing machineries of their hosts, which may explain why PVX P25 affects NbAGO2, having evolved with *Solanaceous* hosts, but not AtAGO2 (Fig. 1).

Our analysis also underlines the usefulness of the PVX-Arabidopsis pathosystem for dissecting the multifactorial nature of virus resistance in plants. Three out of forty-seven *JAX1* (-) accessions possess Col-0-like *AGO2* alleles, but were susceptible to PVX, whereas one C24-like accession was not infected by PVX (Fig. 3, Supplementary Table 1). Likewise accessions Stepn-2 and Bolin-1 are hypersusceptible to PVX (Fig. 3a). These examples suggest that there are likely additional genetic factors that contribute to susceptibility and resistance to PVX (and potentially other viruses) beyond that provided by AGO2 variants.

Both Col-0-like and C24-like alleles have remained prevalent in Eurasian populations at near equal frequencies (Fig. 2). Balancing selection on alleles conferring differing degrees of disease resistance is thought to occur because of trade-offs between fitness and defense under differing environmental conditions^40,41^. Indeed, Col-0-like AGO2 variants may confer greater virus resistance but have a negative effect on plant fitness. Alternatively, it is possible that C24-like *AGO2* alleles are more functional against other naturally-occurring viruses in *Arabidopsis* or that it confers some other fitness advantage unrelated to its role in antiviral defense. Such conservation of a non-functional allele of a gene involved in antiviral silencing has been described for the *RDR1* gene in *N. benthamiana*^13^ wherein a functional antiviral allele negatively impacts early vigor in *N. benthamiana*^13^. Likewise, it could be speculated the frequent occurrence of non-functional *JAX1* alleles^19^ (Fig. 3, Supplementary Table 1) may be due to a fitness penalty for this type of resistance to potexviruses. However, AGO2 differs from these examples in that the C24-like variants retain most of their inherent activities (Fig. 5). Interestingly, it has been reported that *AGO2* has undergone positive selection in tomato during the domestication process^42^, although what function has been selected for is unclear.

The most frequent polymorphisms in the *AGO2* coding sequence, ΔGR and D33G, are both found in a region of the protein identified as Block 43, (Fig. 2b, Fig. 3)^43^. This motif is present in different copy numbers in different AGO proteins from animal and plant species and, although well conserved, its function is unknown^43^. It is of notable that in nearly all cases these two polymorphisms are either both absent or both present (Fig. 2), suggesting that their pairing may have some functional significance. Indeed, many C24-like variants have different GR deletions, suggesting that they may have arisen independently (Supplementary Fig. 3), possibly to compensate for a loss of function due to having an aspartate at residue 33. Such a recurrent selection for alleles non-functional for disease resistance would be similar to selection at certain NLR loci, such as RPM1, where multiple loss of function alleles have been selected for over evolutionary time^44,45^.

Among *Arabidopsis* AGO proteins, AGO1, 2, 3, 5 and 10 encode multiple B43 motifs^43^. In numerous animal systems and more recently in *Arabidopsis*, arginine residue within this block have been shown to be methylated by PMRT^46–51^. In *Arabidopsis*, bacterial infection results in a reduction in expression of *PMRT5*, which in turn reduces arginine methylation of AGO2, leading to stabilization of the protein and miRNA loading^51^. Consistent with this, we also observe greater accumulation of AGO2 in both Col-0 and C24 during bacterial infection (Fig. 5b). Col-0-like AGO2 proteins possess more GR repeats than C24-like AGO2 proteins, which could potentially be modified by PMRT5 activity. However, since both Col-0 and C24 appear to accumulate to similar levels (Fig. 4, Fig. 5, Fig. 6), and because both AGO2 proteins have similar functions in antibacterial defense and DNA methylation (Fig. 4), we deem it unlikely that these polymorphisms impair protein stability or sRNA loading. Alternatively, we suggest that the differences may prevent the P25 VSR activity from inhibiting Col-0-AGO2, although this will require further study.

### Conclusion

We demonstrate that natural variation in a core RNA silencing protein, between and within species, is a major factor determining susceptibility to PVX. These results have important implications for plant-virus coevolution and suggest that identification of natural variants in *AGO2* in crop and related species may prove useful in developing crops with increased resistance to virus pathogens.

## Methods

### Plant material and growth conditions

*Nicotiana benthamiana* and *Arabidopsis thaliana* plants were grown in soil (BM6, Berger and Agromix, Fafard PLACE, respectively) in growth chambers with 16-h-light/8-h-dark and 12-h-light/8-h-dark photoperiod at 23°C and 21°C respectively. Col-0 (CS28168), C24 (CS28127) and *Arabidopsis* wild accessions (CS76427) were obtained from the ABRC stock center. The *ago2-1* mutant in Col-0 background has been described elsewhere^52^. RILs between Col-0 and C24 were kindly provided by R.C. Meyer and have been described previously^20^.

### Plasmid construction and transient expression

The *NbAGO2* ORF was cloned using cDNA derived from TBSV infected *N. benthamiana* leaves using primers listed in Supplementary Table 4. For the generation of different *Arabidopsis* AGO2 expression clones, cDNA (Fig. 1) and gDNAs (Fig. 6, Supplementary Fig. 6 and Supplementary Fig. 7) of the appropriate accessions were used as templates for PCR amplification with primers listed in Supplementary Table 4. PCR products were purified and cloned into the pGEM-T easy vector (Promega) and subcloned into the XbaI and BamHI sites of pBIN61 vector containing an N-terminal FLAG epitope in frame with the XbaI site or in pBIN61 empty vector for HA-tagged NbAGO2. All other constructs have been previously described including PVX, PVX-GFP, PVX-GFPΔTGB and PlAMV-GFP binary constructs^19,53–55^, as well as FLAG-P25^15^ and pBIC-HA-AtAGO2^52^. For the generation of an AGO2 expression vector under the control of the *NbAGO2* promotor, AGO2 upstream regulatory sequences (2068 bp) were amplified from genomic DNA with primers listed in Supplementary Table 4. PCR products were purified and cloned into the pGEM-T easy vector (Promega) and subcloned into the Acc65I and XbaI sites sites of pBIN61 to replace the 35S promoter. *Agrobacterium*-mediated transient expression (agroinfiltration) assays in *N. benthamiana* were performed as previously described^15^.

### Virus inoculation

Infections of three-week-old *Arabidopsis thaliana* plants were carried out by rub inoculation as previously described^15^. Briefly, saps were produced with PVX-infected *N. benthamiana* plant material. Infected material was ground in 0.1 M phosphate buffer, pH 7.0 (2 mL/g of infected tissue). Mock inoculations were performed with sap produced with uninfected *N. benthamiana* plants (2 mL/g of healthy tissues).

### Protein extraction and analysis

Protein extraction and analysis was carried out as previously described^15^. Proteins were detected by immune blotting using anti-HA-horseradish peroxidase conjugated (HRP) antibodies (Sigma, 1:3,000 dilution), anti-FLAG-HRP antibodies (Sigma, 1:5,000 dilution), anti-GFP-HRP antibodies (Santa Cruz, 1:3,000 dilution), anti-PVX-CP rabbit polyclonal antibodies (Agdia, 1:3,000 dilution) and anti-AGO2 antibody (Agrisera, 1:5,000 dilution or Abiocode 1:7,500). Detection of the latter three primary antibodies was performed using donkey anti-IgG rabbit-HRP polyclonal antibodies (BioLegend, 1:10,000 dilution).

### Gene expression and DNA methylation analysis

Total RNA was isolated with Trizol (Ambion) and subjected to RT-PCR using primers listed in Supplementary Table 4. Gene expression and McrBC analyses were performed as previously described^27^ with minor modifications. 10 µg of genomic DNA were digested with 6u of McrBC (NEB) in a final volume of 50 µl for 3h at 37°C. The digested DNA was then analyzed by semi-quantitative PCR using primers listed in Supplementary Table 4.

RT-qPCR was performed as previously described^16,56^ using primers listed in Supplementary table 4.

### Quantification of SA

Quantification of SA was performed by PhenoSwitch Bioscience Inc. (Sherbrooke, Canada). Briefly, salicylic acid was extracted from crushed tissues by the addition of 500µl of methanol containing 0.01 ng/µl of internal standard (D4-Salicylic acid). The samples were then incubated at 4°C for 30 minutes with end-over-end mixing and the insoluble material was cleared by centrifugation. The supernatant was diluted 10 fold in water and the pH was adjusted to 7 by the addition of 50 mM ammonium acetate. A standard curve containing 500, 250, 125, 62.5, 31.25, 15.62, 7.81, 3.91, 1.95 or 0.97 µg/ml SA was prepared in 50 µl H2O and processed the same way as the plant samples. For weak anion exchange solid phase extraction (WAX SPE) of salicylic acid and internal standard, the protocol was as follows: phase conditioning with acetonitrile, wash with 50 mM ammonium acetate, sample loading, wash with 50 mM ammonium acetate, elution with 5% ammonium hydroxide in water. The eluate was dried down by speed vac, reconstituted in 50µl of water containing 0.2% formic acid and 10 mM ammonium formate and processed by LC-MS/MS.

## Acknowledgements

We are grateful to Rhonda C. Meyer for the RILs (C24 x Col-0) collection. This work was supported by funding from the National Science and Engineering Council (Canada) and the Fonds de Recherche du Québec, Nature et Technologie (FRQNT) to P.M., by a scholarship from the Chinese Scholarship Council to Z.Z. and by a graduate fellowship from the NSERC CREATE Agrophytosciences program to A.A and C.R.L..

## Contributions

C.B. and P.M. conceived and designed the experiments. C.B., A.A., C.R.L., Z.Z. and S.B. performed the experiments. C.B. and A.A. analyzed the data. C.B. and P.M. wrote the article.

## Competing interests

The authors declare no competing interests.

**Supplementary Figure 1.**
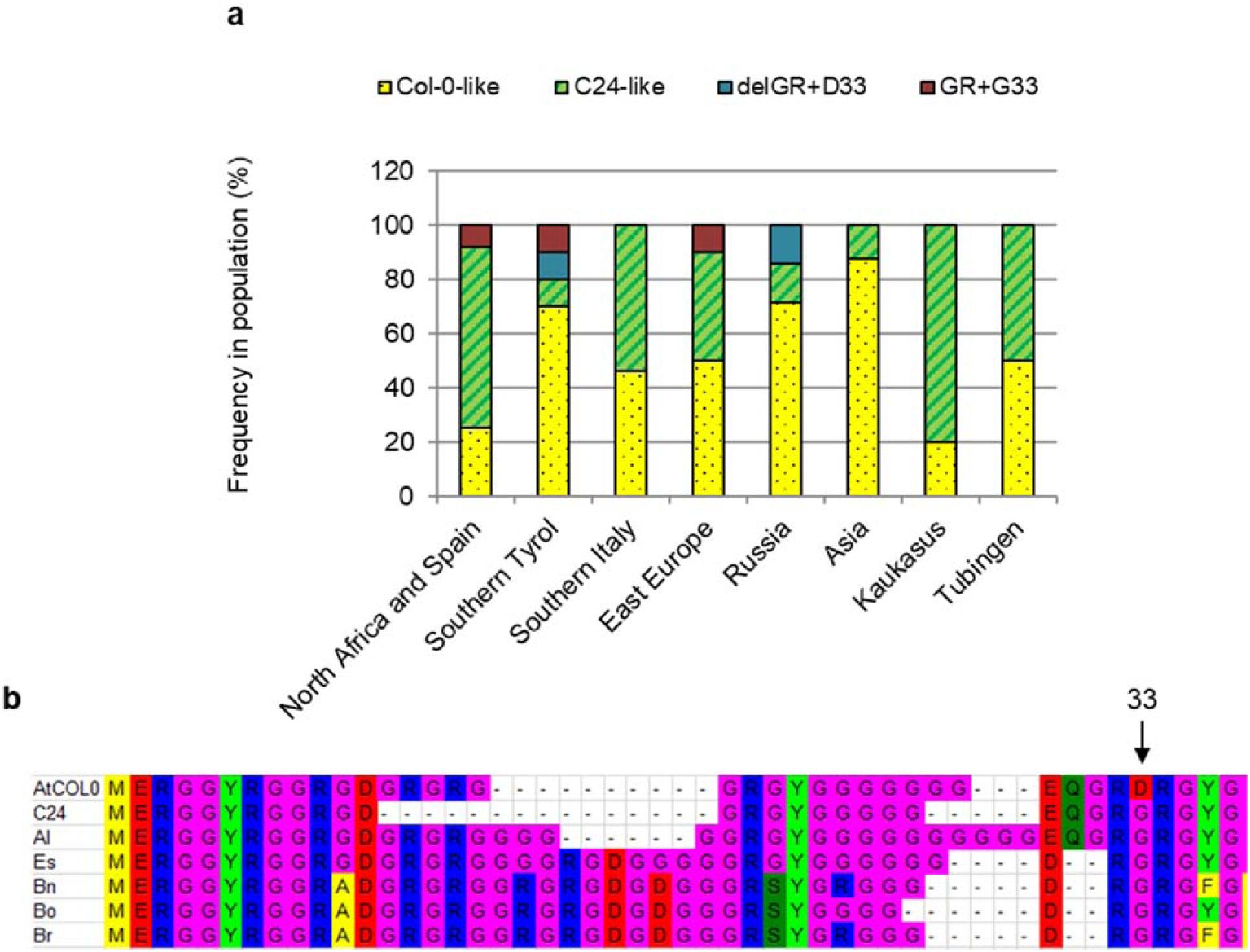
C24 and Col-0 alleles are found in all eight Eurasian populations analyzed. **a**, Chart representing allele frequencies in eight different Eurasian populations. Note that alleles have been classified based only on polymorphisms found in the GR motif and residue 33 of *AGO2* regardless of polymorphisms found elsewhere in the coding sequence. **b**, Amino acid alignment of a portion of the N-terminus of AGO2 from *Arabidopsis thaliana* Col-0 (AtCol-0), C24 (At-C24) and *Arabidopsis lyrata* (Al), as well as other *Brassicaceae, Eutrema salsugineum* (Es), *Brassica napus* (Bn), *Brassica oleracea* (Bo) and *Brassica rapa* (Br).

**Supplementary Figure 2.**
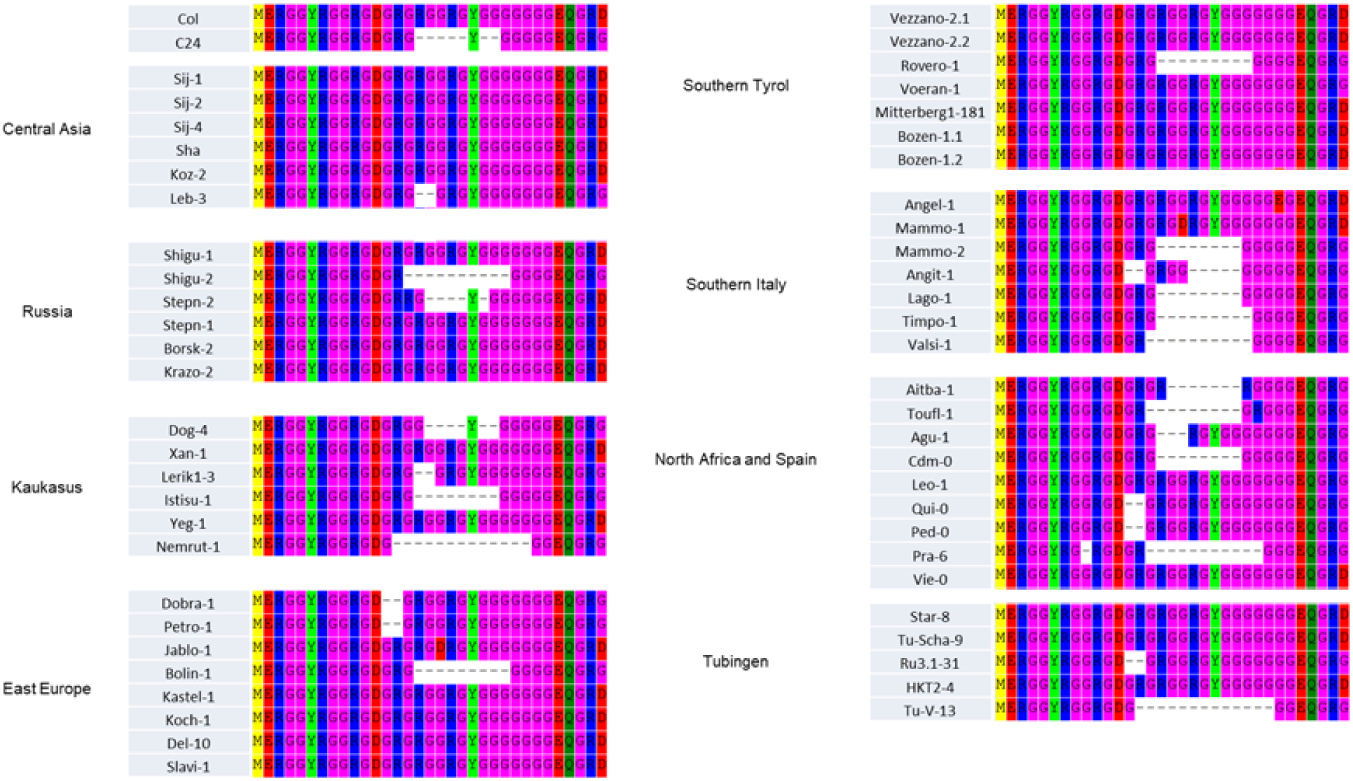
Alignment of the N-terminus of AGO2 of accessions tested in Figure 3. The first thirty-seven residue of Col-0 AGO2 were aligned using ClustalW with manual adjustments.

**Supplementary Figure 3.**
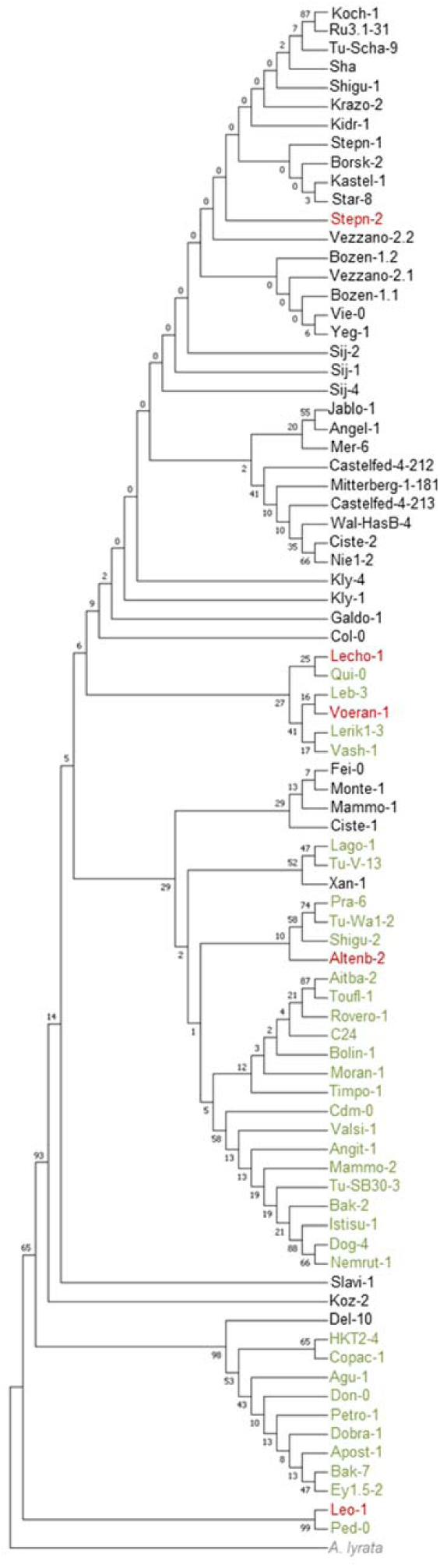
Phylogenetic analysis of Arabidopsis AGO2 sequences. Phylogenetic analysis was conducted on the entire Eurasian Arabidopsis accessions dataset and include Col-0 and C24 as well as *Arabidopsis lyrata AGO2* as an outgroup. Sequences were aligned with ClustalW, using the following alignment parameters: for pairwise alignment, gap opening, 10.0, and gap extension, 0.1; for multiple alignment, gap opening, 10.0, and gap extension, 0.20. Resulting alignments were analysed using Molecular Evolutionary Genetics Analysis 5 (MEGA5) software to generate a neighbor-joining tree derived from 5000 replicates. The evolutionary distances were computed using the JTT matrix-based method [3] and are in the units of the number of amino acid substitutions per site. The percentage of replicate trees in which the associated taxa clustered together in the bootstrap test (5000 replicates) are shown next to the branches. Accession names are colored according to their *AGO2* allele: Col-0-like are black, C24-like are green and rare alleles are red.

**Supplementary Figure 4.**
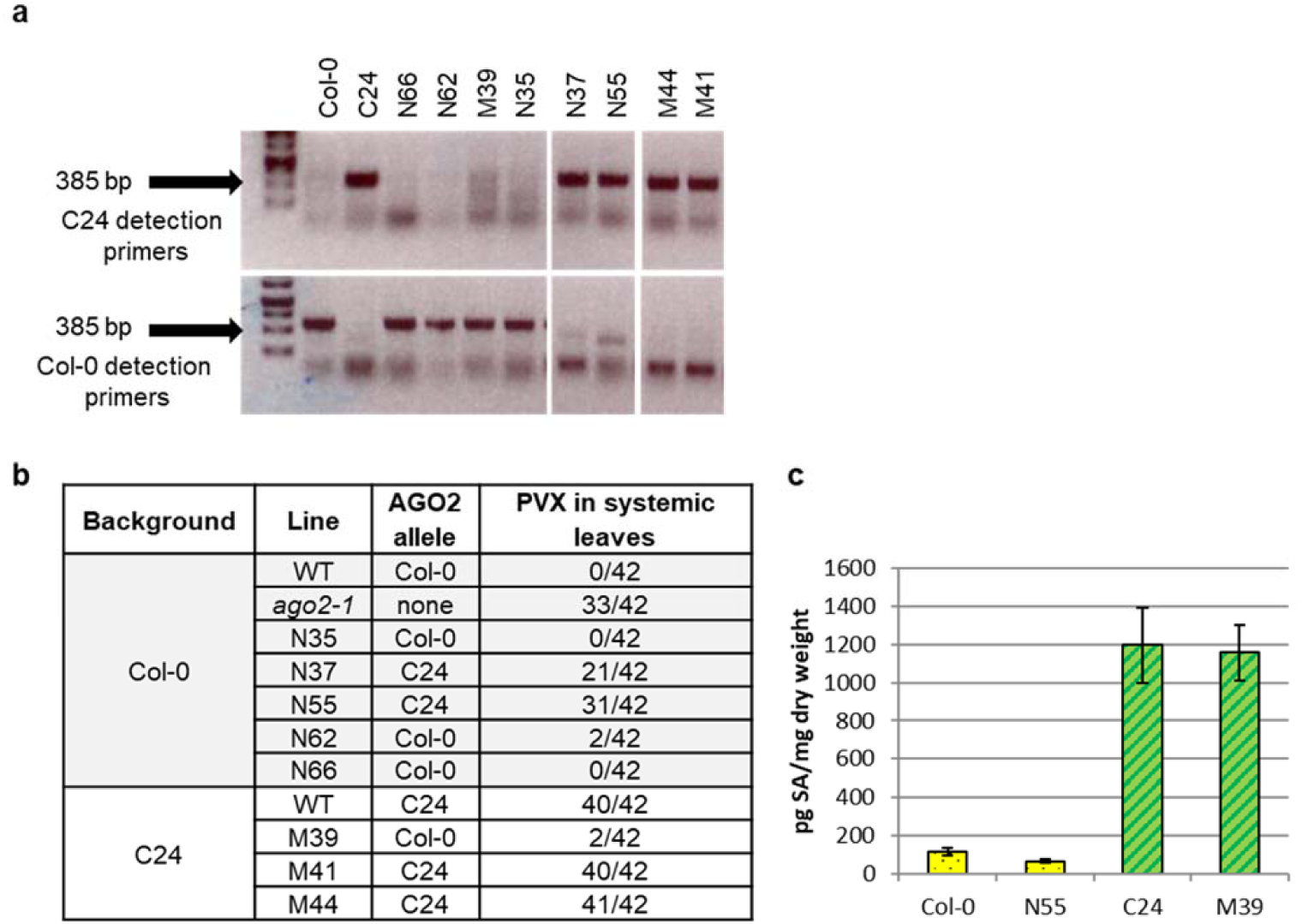
Correlation between *AGO2* alleles in RILS and susceptibility of *Arabidopsis* to PVX. **a**, Verification of *AGO2* alleles by PCR on gDNA of RILs with allele-specific primers. **b**, Numbers of PVX-inoculated plants showing systemic infection were scored by anti-PVX CP immunoblotting at 21 dpi. **c**, Quantification of SA content by LC-MS/MS in two different RILs and their respective background accessions, namely Col-0 and C24.

**Supplementary Figure 5.**
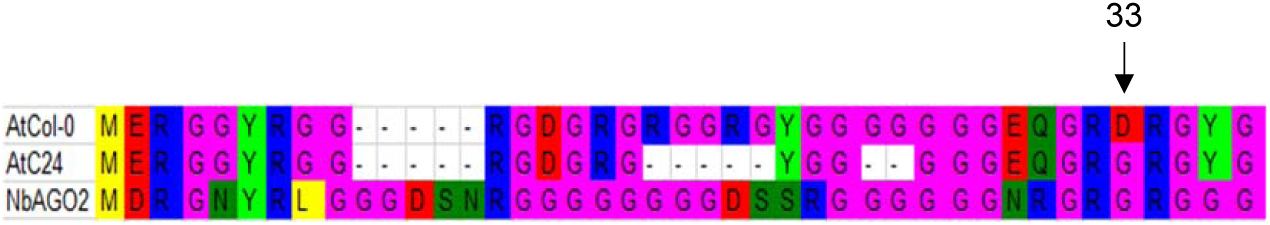
Alignment of the N-termini of AGO2 proteins tested. The first thirty-seven residues of Col-0 AGO2 were aligned with *Nicotiana benthamiana* (NbAGO2) and *Arabidopsis* C24 (C24) AGO2 sequences using ClustalW with manual adjustment.

**Supplementary Figure 6.**
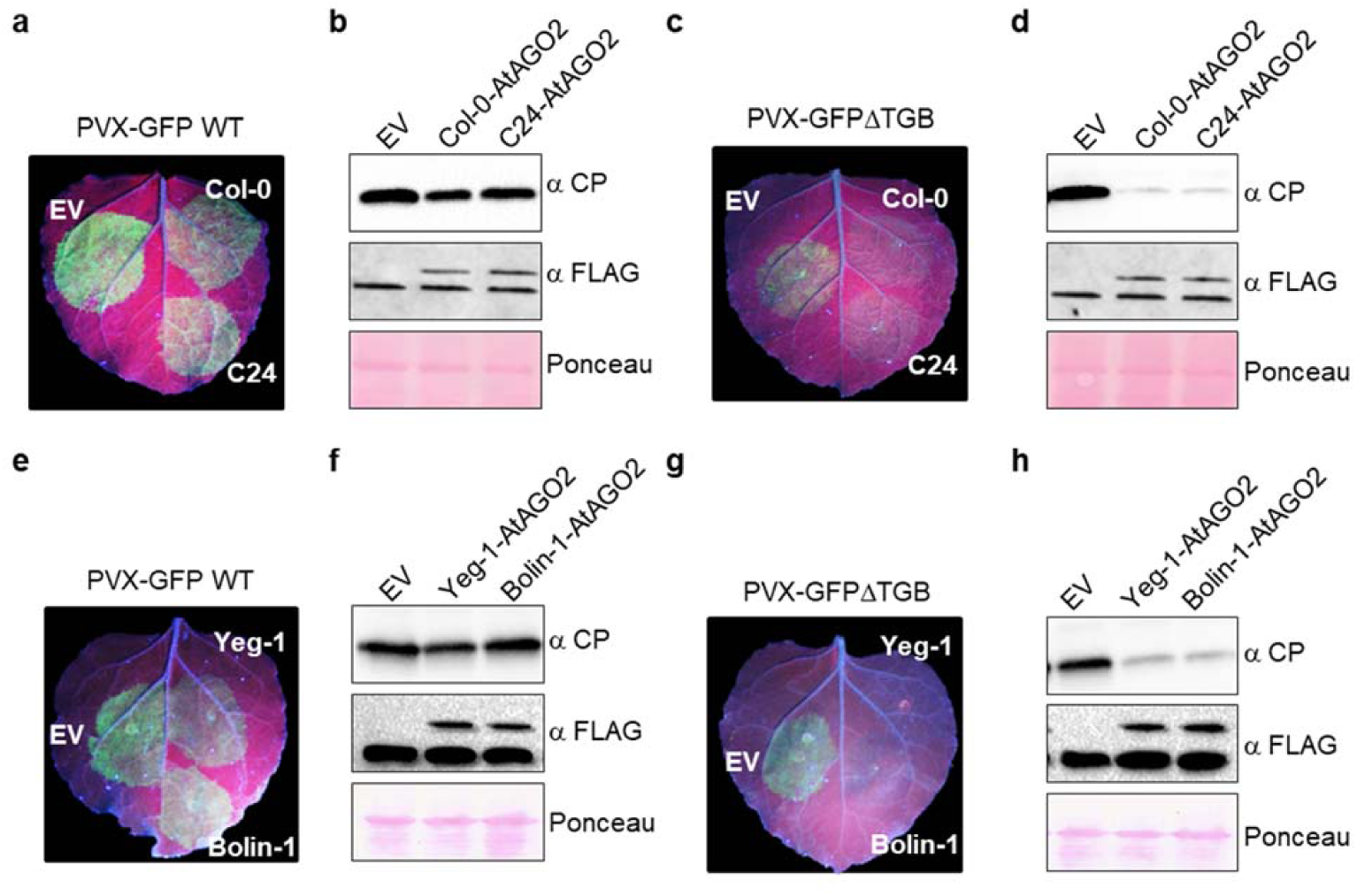
Polymorphisms found in C24 and C24-like *AGO2* affect its antiviral activity in *N. benthamiana*. **a**, **c**, **e**, **g**, *N. benthamiana* leaves were agroinfiltrated with PVX-GFP WT **a** and **c** or ΔTGB **e** and **g**, along with 35S:FLAG-Col-0-AGO2, 35S:FLAG-C24-AGO2 or empty vector (EV) **a** and **c** or with 35S:FLAG-AtYeg-1-AGO2, 35S:FLAG-AtBolin-1-AGO2 or empty vector (EV) **e** and **g**. Leaves were photographed under UV illumination at 4 dpi. **b**, **d**, **f**, **h**, Total protein extracts were prepared from *N. benthamiana* leaves agroinfiltrated as in **a**, **c**, **e**, **g** at 4 dpi and subjected to SDS-PAGE, followed by anti-PVX CP (top panel) or anti-FLAG (middle panel) immune blotting. Ponceau staining (bottom panel) of the same extracts is shown to demonstrate equal loading.

**Supplementary Figure 7.**
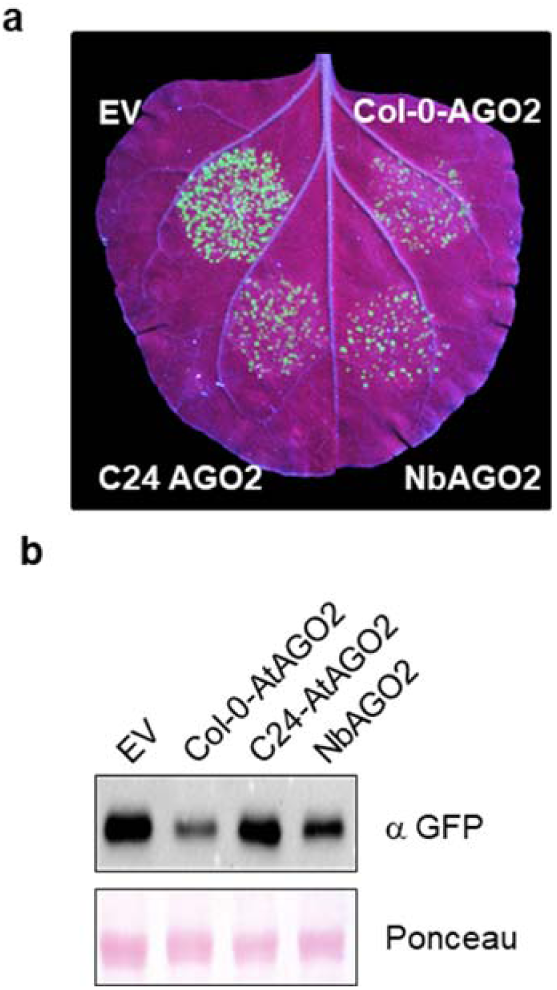
Differential antiviral activity of AGO2 variants against PlAMV. **a**, *N. benthamiana* leaves were agroinfiltrated with PlAMV-GFP along with 35S:FLAG-AtCol-0-AGO2, 35S:FLAG-AtC24-AGO2, 35S:HA-NbAGO2 or empty vector (EV). Leaves were photographed under UV illumination at 4 days post infiltration (dpi). **b**, Total protein extracts were prepared from *N. benthamiana* leaves agroinfiltrated as in **a** at 4 dpi and subjected to SDS-PAGE, followed by anti-GFP immunoblotting (top panel). Ponceau staining (bottom panel) of the same extracts is shown to demonstrate equal loading.

